# Disambiguating the role of blood flow and global signal with Partial Information Decomposition

**DOI:** 10.1101/596247

**Authors:** Nigel Colenbier, Frederik Van de Steen, Lucina Q. Uddin, Russell A. Poldrack, Vince D. Calhoun, Daniele Marinazzo

## Abstract

In resting state functional magnetic resonance imaging (rs-fMRI) a common strategy to reduce the impact of physiological noise and other artifacts on the data is to regress out the global signal using global signal regression (GSR). Yet, GSR is one of the most controversial preprocessing techniques for rs-fMRI. It effectively removes non-neuronal artifacts, but at the same time it alters correlational patterns in unpredicted ways. Furthermore the global signal includes neural BOLD signal by construction, and is consequently related to neural and behavioral function. Performing GSR taking into account the underlying physiology (mainly the blood arrival time) has been proved to be beneficial. From these observations we aimed to: 1) characterize the effect of GSR on network-level functional connectivity in a large dataset; 2) assess the complementary role of global signal and vessels; and 3) use the framework of partial information decomposition to further look into the joint dynamics of the global signal and vessels, and their respective influence on the dynamics of cortical areas. We observe that GSR affects intrinsic connectivity networks in the connectome in a non-uniform way. Furthermore, by estimating the predictive information of blood flow and the global signal using partial information decomposition, we observe that both signals are present in different amounts across intrinsic connectivity networks. Simulations showed that differences in blood arrival time can largely explain this phenomenon. With these results we confirm network-specific effects of GSR and the importance of taking blood flow into account for improve denoising methods. Using GSR but not correcting for blood flow might selectively introduce physiological artifacts across intrinsic connectivity networks that distort the functional connectivity estimates.

## Introduction

In recent years there has been increasing interest in the use of resting state functional magnetic resonance imaging (rs-fMRI) in neuroimaging research. A popular approach in rs-fMRI is to map the functional architecture of the human brain using patterns in resting-state or intrinsic correlations (Michael D. Fox & Raichle, 2007). The correlations of low frequency oscillations present in the blood oxygenation level dependent (BOLD) signal reflect the functional connectivity between different brain regions. Regions that then show a high mutual correlation are referred to as a resting-state network (RSN) or intrinsic connectivity network (ICN). In rs-fMRI studies a common pre-processing step before estimating functional connectivity is global signal regression (GSR), where a global time course is regressed out of the data. The global signal (GS) is obtained by averaging the resting-state time courses over the entire brain (Desjardins, Kiehl, & Liddle, 2001). The GS is often thought of as a mixture of processes that confound the BOLD fMRI signals (artifacts). Based on the assumption that processes that are globally spread across the brain cannot be linked to neuronal activation, it could be beneficial to remove them to denoise the data. Indeed, fluctuations in the GS have been linked to physiological fluctuations, mainly respiratory effects, head motion, hardware scanner related effects and vascular effects (Murphy & Fox, 2017; Power, Plitt, Laumann, & Martin, 2017).

However, in addition to these non-neuronal confounds, GSR has also been related to neuronal fluctuations. Schölvinck and colleagues showed that the fMRI BOLD signal calculated over the entire cerebral cortex in monkeys showed positive correlations with the spontaneous fluctuations in the local field potentials in a single cortical site (Schölvinck, Maier, Ye, Duyn, & Leopold, 2010). Others have shown a relationship between the global signal and EEG measures of vigilance and broadband electrical activity using simultaneous EEG and fMRI (Wen & Liu, 2016; Wong, Olafsson, Tal, & Liu, 2013). These issues are well summarized in (T. T. Liu, Nalci, & Falahpour, 2017) which set to disambiguate the nuisance and the information component of the GS by looking at its origin and associations with other neuroimaging measures. More recent studies have demonstrated that the activity of the basal forebrain is intimately linked to cortical arousal (X. Liu et al., 2018) and GS (Turchi et al., 2018), showing that inactivation of the basal forebrain leads to increased global spontaneous fluctuations. The GS has been explicitly connected to these fluctuations, either by predicting the nature of the quasiperiodic patterns of large-scale brain activity (Yousefi, Shin, Schumacher, & Keilholz, 2018), or by encoding the transitions between them (Gutierrez-Barragan, Basson, Panzeri, & Gozzi, 2018). Using calcium based imaging in mice Matsui and colleagues identified global propagating waves of activity in the neocortex of mice, which points to the existence of global neuronal signals (Matsui, Murakami, & Ohki, 2016). In sum, evidence demonstrates that the GS is a mixture of neuronal and non-neuronal components, but it’s unclear how much each component contributes to the GS (Uddin, 2017). As a result, GSR may remove fluctuations of neuronal origin, which could induce errors in functional connectivity estimates (Chen et al., 2012). After all, the whole brain can be thought as the roughest parcellation in terms of ROIs or Intrinsic Connectivity Networks, and in this sense it contains all neural/behavioral correlates, as well as all the nuisance.

The practice of GSR has been debated since Murphy and colleagues disputed the finding of anti-correlations between the default-mode network (DMN) and the task positive network (TPN) reported by Fox and colleagues (M. D. Fox et al., 2005; Murphy, Birn, Handwerker, Jones, & Bandettini, 2009). By using a mathematical argument Murphy and colleagues showed that the anti-correlations found were an artifact introduced by GSR, and in the absence of this preprocessing step the DMN and TPN were positively correlated. Mathematically GSR shifts the distribution of functional connectivity estimates to be centered around zero, thereby inducing the existence of both positive and negative correlations (Murphy et al., 2009). Similarly, other studies also observed anti-correlations only when GSR was applied (Ibinson et al., 2015; Weissenbacher et al., 2009), while others have found anti-correlations without applying GSR, but after ingestion of caffeine (Wen & Liu, 2016; Wong et al., 2013), physiological noise correction (Chang & Glover, 2009) or component-based noise reduction (Chai, Castañón, Öngür, & Whitfield-Gabrieli. 2012), suggesting that anti-correlations might not be an artifact induced by GSR and have some neuronal origin. The nature of the anti-correlations as a true reflection of functional connectivity versus an artifact of the GSR technique is not clear nor agreed upon. But most evidence seems to point to a reduction in positive correlations and an increase in spurious anti-correlations. Moreover, using a modeling approach it has been shown that GSR does more than “just” inducing anti-correlations, also altering the underlying correlation structure in unpredictable ways. Saad and colleagues showed in a group comparison study that applying GSR alters short-and long-range correlations within a group, leading to spurious group differences in regions that were not modeled to show true functional connectivity differences (Saad et al., 2012).

Despite the critiques a consensus on the use of GSR has not been reached, and it remains a popular denoising method in rs-fMRI studies (Murphy & Fox, 2017; Power, Plitt, et al., 2017). A recent wave of studies further proves its usefulness in reducing the impact of artifacts, particularly those resulting from participant head motion. Studies comparing the effectiveness of different denoising approaches show that only combinations of preprocessing pipelines that include GSR can minimize the effects of motion (Ciric et al., 2017: Lydon-Staley, Ciric, Satterthwaite, & Bassett, 2018: Parkes, Fulcher, Yücel, & Fornito, 2018; Satterthwaite et al., 2017), temporally lagged artifacts (Byrge & Kennedy, 2018) and respiration (Power, Laumann, Plitt, Martin, & Petersen, 2017).

Recent work by Erdoğan and colleagues proposed an improvement to the standard GSR method called dynamic GSR (dGSR) which improves the sensitivity of functional connectivity estimates (Erdoğan, Tong, Hocke, Lindsey, & deB Frederick, 2016). dGSR effectively deals with systemic low frequency oscillations (sLFO’s) that are non-neuronal of origin and oscillate at frequencies of interest in rs-fMRI (−0.01-0.1 Hz). Here we refer to sLFO as fluctuations of clear non-neuronal origin (as defined in the cited studies), as opposed to quasiperiodic patterns (QPPs) which reflect neural activity. These intrinsic signals appear to have a vascular origin that travel with the blood through the body (Tong & Frederick, 2012; Tong et al., 2013) and brain (Tong & Frederick, 2010). The sLFO’s appear in the arteries before propagating through the brain with the cerebral blood flow from large arteries to large veins (Tong, Yao, Jean Chen, & deB Frederick, 2018). The spatial-temporal pattern of the arrival time of these sLFO’s arc consistent with blood flow circulation patterns obtained with dynamic susceptibility contrast (DSC) bolus track imaging (Tong et al., 2017). The method is based on taking the temporal information of blood arrival time into account. For each voxel an optimally delayed version of the global signal is regressed out, which effectively removes the systemic noise of these sLFO’s. As a result, the strength and specificity of functional connectivity estimates is improved compared with standard GSR. In addition, in further research they show that BOLD signal from major arteries (internal carotid artery) and prominent veins (superior sagittal sinus, SSS and internal jugular vein, great vein of Galen) are highly correlated with the GS (T. T. Liu et al., 2017; Tong, Yao, Jean Chen, et al., 2018), supporting the importance of a vascular component in the GS. From this group of studies, it’s evident that a large portion of the GS has a macro-vascular origin, and that considering the temporal blood flow information improves de-noising methods. Indeed, taking information from the vessels in de-noising approaches is important, as recent work has shown that the dynamic effects of vessel information has complex consequences on BOLD responses (Kay et al., 2019). Some recent efforts have been made to merge perfusion and BOLD to take blood into account in functional connectivity studies (Tak, Polimeni, Wang, Yan, & Chen, 2015; Cohen, Nencka, Lebel, & Wang, 2017).

In a recent correspondence, three directions forward are suggested to further evaluate the use of GSR (Power, Laumann, et al., 2017; Power, Plitt, et al., 2017: Uddin, 2017). One of these directions raises the concern that the field lacks empirical studies that focus on the effect of GSR in high dimensional fMRI data. In this study, we attempt to address this issue by applying GSR to high dimensional empirical fMRI data, using an information theory and correlation approach to investigate how functional connectivity estimates are affected by its use. We are interested in a more detailed evaluation of which brain regions and intrinsic connectivity networks in the connectome are affected by GSR. In line with the work from (Gotts et al., 2013; Saad et al., 2012), we expect that the large scale brain connectivity pattern will be altered by GSR in a regional and network-specific way.

Furthermore, from the work of (He, Shin, & Liu, 2010; T. T. Liu et al., 2017; Tong, Yao, Chen, & Frederick, 2018) we know that the GS and vessel BOLD signals are highly related, and affect the statistical dependencies in the connectome (Erdoğan et al., 2016). We set out to investigate their unique and joint informative role in the connectome, beyond this high correlation. By applying multiscale partial decomposition of predictive information (Faes, Marinazzo, & Stramaglia, 2017) we were able to disambiguate the reduction of uncertainty due to the GS and vessel BOLD signal across intrinsic connectivity networks and multiple time scales. For the first time, we apply an information theoretical approach to the debate of GS that goes beyond a correlational framework. This could further help us clarify the effects of GSR, and the information we get out of the BOLD signal at different spatiotemporal scales.

In short, the main goals of this study are: 1) characterizing the effect of GSR on network-level functional connectivity estimates in a large dataset; 2) assess the complementary role of global signal and vessels BOLD signal in this modulation; 3) use the framework of (predictive) information decomposition to further examine these joint dynamics. We use a large public dataset of rs-fMRI data, simulations and simultaneous calcium and hemodynamic recordings from mouse data. We would like to emphasize that to the goal is to complement the existing literature without explicitly advocating for or against the use of GSR.

## Materials and Methods

### HCP rs-fMRI data

We used the publicly available 900 Subject release data of the Human Connectome Project (HCP; Van Essen et al., 2012). The full 900 Subject release contains MRI data from 899 healthy participants. The rs-fMRI data from 34 subjects were excluded during the GSR analyses, and another 99 subjects during calculation of the partial information decomposition (PID) due to technical problems during processing on our high-performance cluster. Therefor rs-fMRI data from 865 subjects were used for the GSR analyses, and data from 766 subject for the Partial Information Decomposition. See (Smith et al., 2013; Van Essen et al., 2012) for details of the acquisition procedures.

#### Data preprocessing

Three preprocessing pipelines were applied to rs-fMRI data: HCP-PID, HCP-NO GSR and HCP-GSR. All three pipelines underwent the following steps: The minimal HCP preprocessing steps which include gradient distortion correction, head motion correction, image distortion correction, spatial transformation to Montreal Neurological Institute (MNI) space and intensity normalization (Glasser et al., 2013). In addition to the minimal preprocessing, several additional preprocessing steps were performed, including spatial smoothing using a Gaussian filter kernel with 4mm full width at half maximum (FWHM), removing linear trends and nuisance regression step. In the nuisance regression step, a linear regression of multiple nuisance regressors was applied. In the HCP-PID and HCP-NO GSR nuisance regressors consisted of: 1) The physiological noise from the white matter signal; 2) cerebrospinal fluid; and 3) the motion parameters. In the HCP-GSR pipeline nuisance regressors consisted of: 1) The physiological noise from the white matter signal; 2) cerebrospinal fluid; 3) the motion parameters and; 4) the global signal (GS) calculated as the average BOLD signal across all voxels in the brain. The difference between the HCP-PID and HCP-NO GSR/HCP-GSR pipelines is that no additional bandpass-filtering (0.008 Hz - 0.1 Hz) was done on the HCP-PID data and the (PID) was calculated from these data.

As a next step, the average vessel BOLD signal was calculated. The average vessel BOLD signal was calculated by averaging the extracted BOLD signal from the vessels. The mask for vessel BOLD signal was created by carefully thresholding the preprocessed T1w/T2w ratio images. Finally, the signals from each pipeline were averaged in 278 regions of interest (ROIs) using the Shen parcellation (Shen, Tokoglu, Papademetris, & Constable, 2013). We chose the Shen parcellation for the following reasons: 1) It is a widely used parcellation of the brain; 2) the nodes are of comparable size, avoiding differential contributions to the GS due to parcel size; 3) It has enough nodes to have complexity at a global and within network level, but also not too many which could cause too much variability across subjects and lose interpretability; 4) it has a clear mapping to the higher order Yeo ICN parcellation. To further localize the results in intrinsic connectivity networks, each of the ROIs was assigned to one of the 9 networks (7 cortical networks, plus subcortical and cerebellum regions) as classified by Yeo (Yeo et al., 2011).

#### Connectivity

Functional connectivity was calculated using a correlational and mutual information approach. In both approaches the time series of the 278 ROIs obtained from both pipelines in the previous step: (1) HCP-NO GSR and (2) HCP-GSR data were used to calculate functional connectivity. The individual connectivity matrices were not tħresholded or binarized, and ordered (columns and rows) according to the 9 networks (7 cortical networks, plus subcortical and cerebellum regions) as proposed by Yeo (Yeo et al., 2011). In a final step, the connectivity matrices were averaged over subjects.

In the correlational approach, Pearson correlation coefficients (MATLAB command corr) were used to evaluate functional connectivity (FC). With the HCP-NO GSR data, FC was calculated between all pairs of ROI time series to obtain a symmetric 278 × 278 connectivity matrix for every subject. A similar procedure was followed to obtain the 278 × 278 connectivity matrix for every subject with the HCP-GSR data. Additionally, the Pearson correlation coefficients between the ROI time series and the GS were calculated. To investigate the contribution of vessel BOLD signal to this relationship, partial correlation coefficients (MATLAB command *partialcorr*) were calculated between the ROI and GS time series conditioned on the vessel BOLD time series.

In the mutual information (MI) approach, MI and conditioned mutual information (CMI) were calculated using a bin-less rank based approach based on copulas to evaluate connectivity (Ince et al., 2017). With the HCP-GSR data, MI was calculated between all pairs of ROI time series to obtain a symmetric 278 × 278 connectivity matrix for every subject. A similar procedure was followed to obtain the 278 × 278 connectivity matrix for every subject with the HCP-GSR data. Additionally, the MI between the ROI time series and the GS were calculated. To investigate the contribution of vessel BOLD signal to this relationship, CMI was calculated between the ROI and GS time series conditioned on the vessel BOLD time series. The MI measures used here are non-normalized, resulting from the subtraction of two entropy terms.

### Calcium and Hemodynamic mouse recordings

Data from the work of Matsui and colleagues were used, provided by the authors upon request (Matsui, Murakami, & Ohki, 2016, 2018b). We used data from one mouse which contained simultaneous recordings of calcium and hemodynamics. This dataset has the advantage of: 1) having a high resolution of the brain and vasculature structures and 2) is less affected by movement due to the mouse being tightly head restraint and induced with a light anesthesia. However, at the same time anesthesia has the possible disadvantage of affecting the neurovascular coupling of the neuronal and hemodynamic activity (Matsui, Murakami, & Ohki, 2018a). As a result, the dynamics of the functional connectivity might be different compared to recordings from mice who arc awake. For a detailed overview of the animal preparation and acquisition of the simultaneous calcium and hemodynamic recordings see (Matsui et al., 2016, 2018b).

#### Data preprocessing

For a detailed overview of preprocessing steps see the data preprocessing section in (Matsui et al., 2016, 2018b). Briefly, the following steps were undertaken: all calcium and hemodynamic images were corrected for possible within-scan motion. The images were then further co-registered, spatially down-sampled by a factor of two and high pass filtered (> 0.01 Hz). After filtering, the time series were normalized by subtracting the mean and dividing by the standard deviation. As a next step, the GS and average vessel time series for hemodynamics and calcium were calculated. The GS was calculated by averaging the time series of all the pixels within the slice. To extract the time series of the vessels a mask was created that only contained pixels of the vessels. The time series were then put through 2 separated pipelines: one with no further preprocessing (no-GSR data), and another with GSR as a further preprocessing step (GSR data; Matsui et al., 2016, 2018b). As a final step, 38 ROIs as described in (Matsui et al., 2016, 2018b), and a vessel mask was used for averaging BOLD time series (Figure 1).

**Figure 1.**
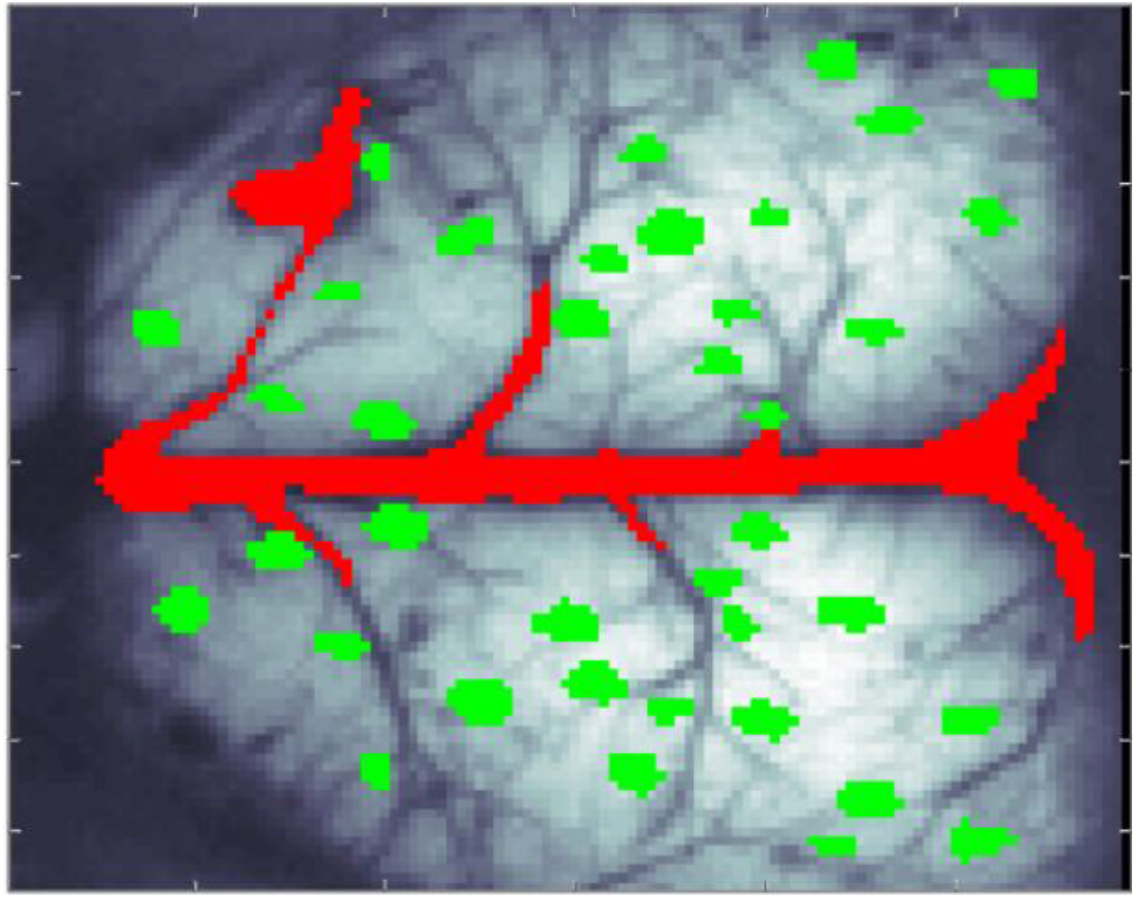
Masks for the Hemodynamic and Calcium mouse recordings. Mask for the 38 ROIs according to Matsui et al., (2016, 2018b) in green, and the mask for the vessel BOLD signal in red.

#### Connectivity

For the correlation approach, FC was calculated between all pairs of ROI time series to obtain a 38 × 38 connectivity matrix for the no GSR data and GSR data. A similar procedure was followed for the mutual information approach, resulting in a 38 × 38 connectivity matrix for the no-GSR and GSR, data.

### Simulations

#### Data generation

For our simulations we used the publicly available simTB toolbox by (Erhardt, Allen, Wei, Eichele, & Calhoun, 2012), which provides flexible generation of fMRI data. In our case we wanted to simulate rs-fMRI BOLD time series that contain a “neuronal” and “blood” contribution. In the simulation setting we considered a resting state design and generated neuronal components with spontaneously generated events over T=1200 time points and TR=0.72 seconds (in line with the HCP data). In total 10 components were generated, 9 neuronal components and one sLFO-carrying blood component. Spontaneously generated event time series from the components were linearly convolved with the canonical hemodynamic response function (HRF; difference of two gamma functions).

Finally, in order to simulate different blood arrival times in each component, we added an linearly increasing delay to the blood component and mixed every neuronal component with a different delayed version of the blood component. In this way, we were able to simulate varying blood arrival time to interacting components. In total, we simulated data for 100 subjects. After simulating the data, GSR was performed on the data resulting in a no-GSR and GSR dataset.

#### Connectivity

For the correlation approach, FC was calculated between all pairs of simulated component time series to obtain a 9 × 9 connectivity matrix for the no-GSR and GSR data. A similar procedure was followed for the mutual information approach, resulting in a 9 × 9 connectivity matrix for the no-GSR and GSR data.

### Partial Information Decomposition

In multivariate systems it makes sense to investigate the joint effect of several driver variables over one (or more) target(s). Partial Information Decomposition (PID) permits calculation of these effects, in particular the unique and joint contributions of two or more driver variables on a target variable. The most relevant measures in this context are the unique information from each of the drivers to the target, and the redundant and synergistic contributions of the two drivers to the target (Figure 2). It is worth noting that Interaction Information, defined as the difference between the mutual information between two variables, and the same quantity conditioned to a third one, also allows definition of redundancy (when it is negative) or synergy (when it is positive). The framework of PID used here provides: 1) distinct non-negative measures of redundancy and synergy, thereby accounting for the possibility that redundancy and synergy may coexist as separate elements of information modification; 2) the disambiguation between drivers and targets.

**Figure 2.**
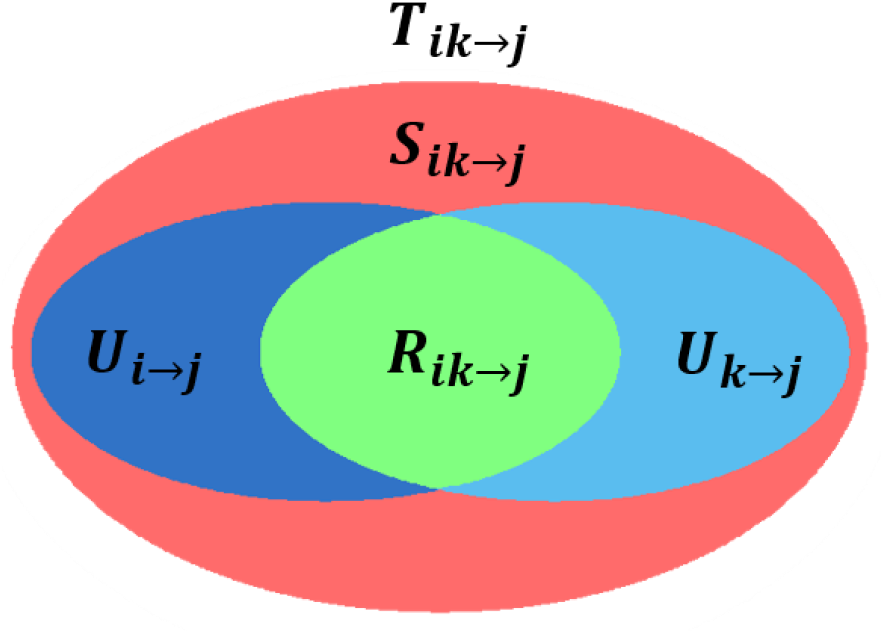
Venn diagram Partial information decomposition (PID). Given two drivers *i* and *k*, and a target *j*, the joint information (T) transferred from the drivers to the target can be decomposed in synergistic (S), redundant (R) and unique (U) contributions.

This framework is valid for several estimators of the joint distributions of the variables. These include the classical information theory measures, in which the distributions are evaluated non-parametrically by binning or other embeddings, and in which information transfer is defined in terms of reduction of uncertainty or surprise, but also simpler estimators in which the distributions are calculated from the joint covariance matrix, and the information transfer has to be interpreted in terms of reduction of joint variance.

Here we use this latter estimator (exact for Gaussian variables, but still robust in the approximate case, and allowing a straightforward extension to the multiscale approach).

PID was performed over multiple temporal scales as described in (Faes et al., 2017).

For the present study, the two drivers are the GS and the BOLD signal extracted from the vessels, and the targets are the timeseries of the individual ROIs. The temporal scale across which we computed PID stretched to about 20 seconds, equivalent to 30-time bins for the human HCP data and the simulated data, 200-time bins for the mouse calcium imaging data, and 100-time bins for the mouse hemodynamics data.

### Data and code availability

The toolbox for the partial information decomposition method is freely available and can be found here: https://github.com/danielemarinazzo/multiscale_PID. The toolbox for Gaussian-Copula mutual information is freely available and can be found here: https://github.com/robince/gcmi. The simTB toolbox we used for simulations can be found here: http://mialab.mrn.org/software/simtb/, and the Matlab script for the simulations and analyses reported in this work can be found here: https://github.com/compneuro-da/GSR_PID.

The HCP dataset we used is publicly available and can be found at https://www.humanconnectome.org/. Some of the figures were made using the Gramm package from Pierre Morel, which can be found at https://github.com/piermorel/gramm and is described in (Morel, 2018).

## Results

### HCP rs-fMRI

#### GSR

A shows the functional connectivity computed using correlations among the 278 ROIs averaged across subjects without (bottom triangle) and with GSR (top triangle) in the HCP dataset. Figure 3B, shows the functional connectivity computed using mutual information for the same dataset with and without GSR. In the correlational approach, GSR reduces functional connectivity across ROI pairs (Figure 3C). We observe that anti-correlations are introduced in the connectome in a non-uniform way across regions and networks (Figure 4). The same observations are made in the mutual information approach, where GSR also reduces the functional connectivity across ROI pairs (Figure 3D), and network-specific effects are found (Figure 5). Two remarks are in order at this point: Mutual information is an unsigned quantities, so the difference cannot be expressed in terms of “negative” correlation induced; there is a nonlinear relation between MI and correlation (see also figure 3 in (Ince et al., 2017), resulting in an enhanced contrast of strong effects with respect to background values.

**Figure 3.**
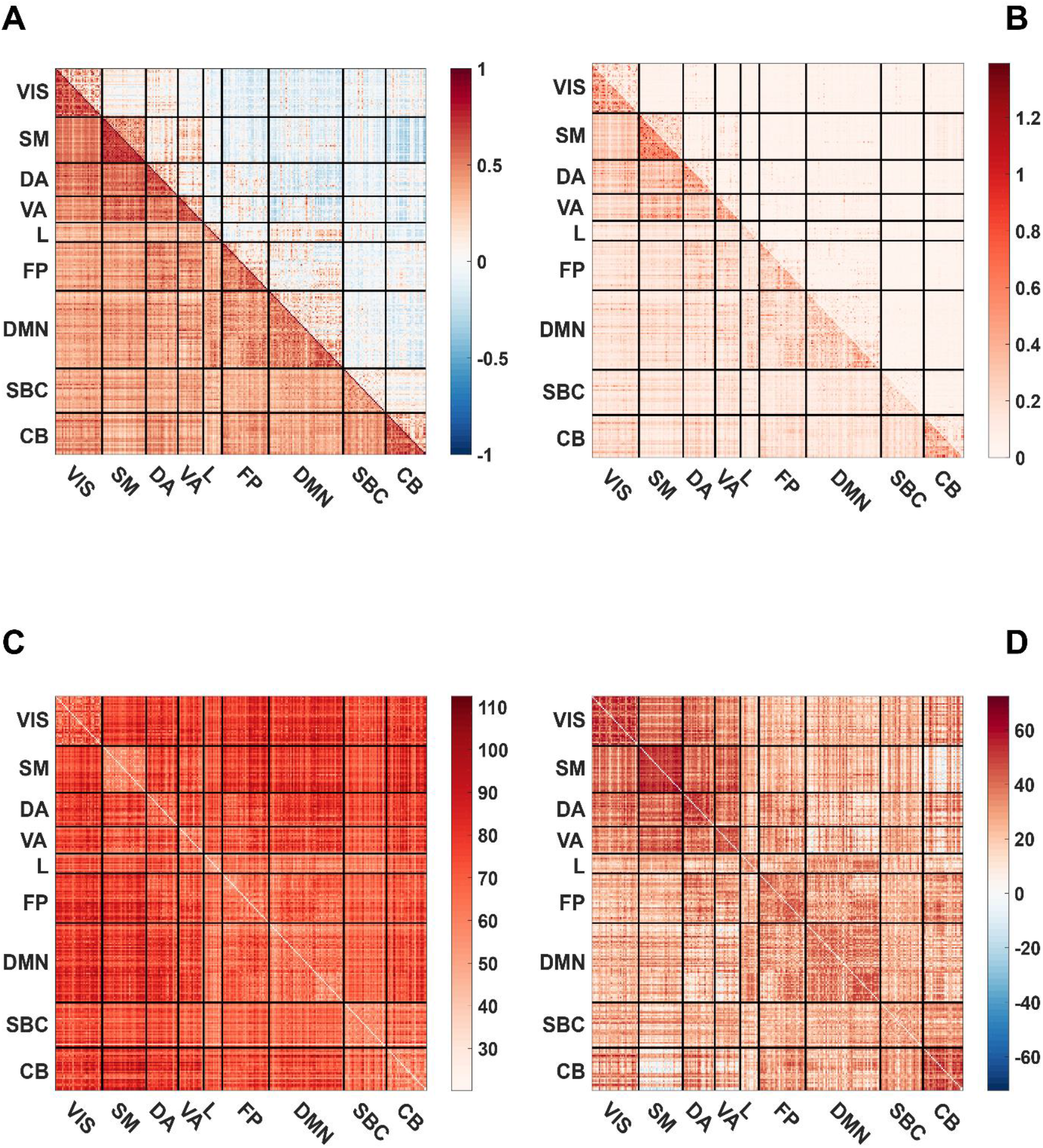
Effect of GSR on functional connectivity in the HCP dataset. ***(A)*** estimated functional connectivity using correlations before (bottom triangle) and after GSR (top triangle). ***(B)*** estimated functional connectivity using mutual information before (bottom triangle) and after GSR (top triangle). ***(C)*** paired *t*-test using permutations between the correlational functional connectivity before and after GSR for every ROI pair. ***(D)*** paired *t*-test using permutations between the mutual information functional connectivity before and after GSR for every ROI pair. For visualization the 278 ROIs are ordered according to the 9 intrinsic connectivity networks of Yeo et al., 2011. The intrinsic connectivity networks arc: VIS=visual network; SM= Somatomotor; DA=Dorsal Attention; VA=Ventral Attention; L=Limbic; FP=Frontoparietal; DMN=Default Mode Network; SBC Subcorlical; CB Cerebellum.

**Figure 4.**
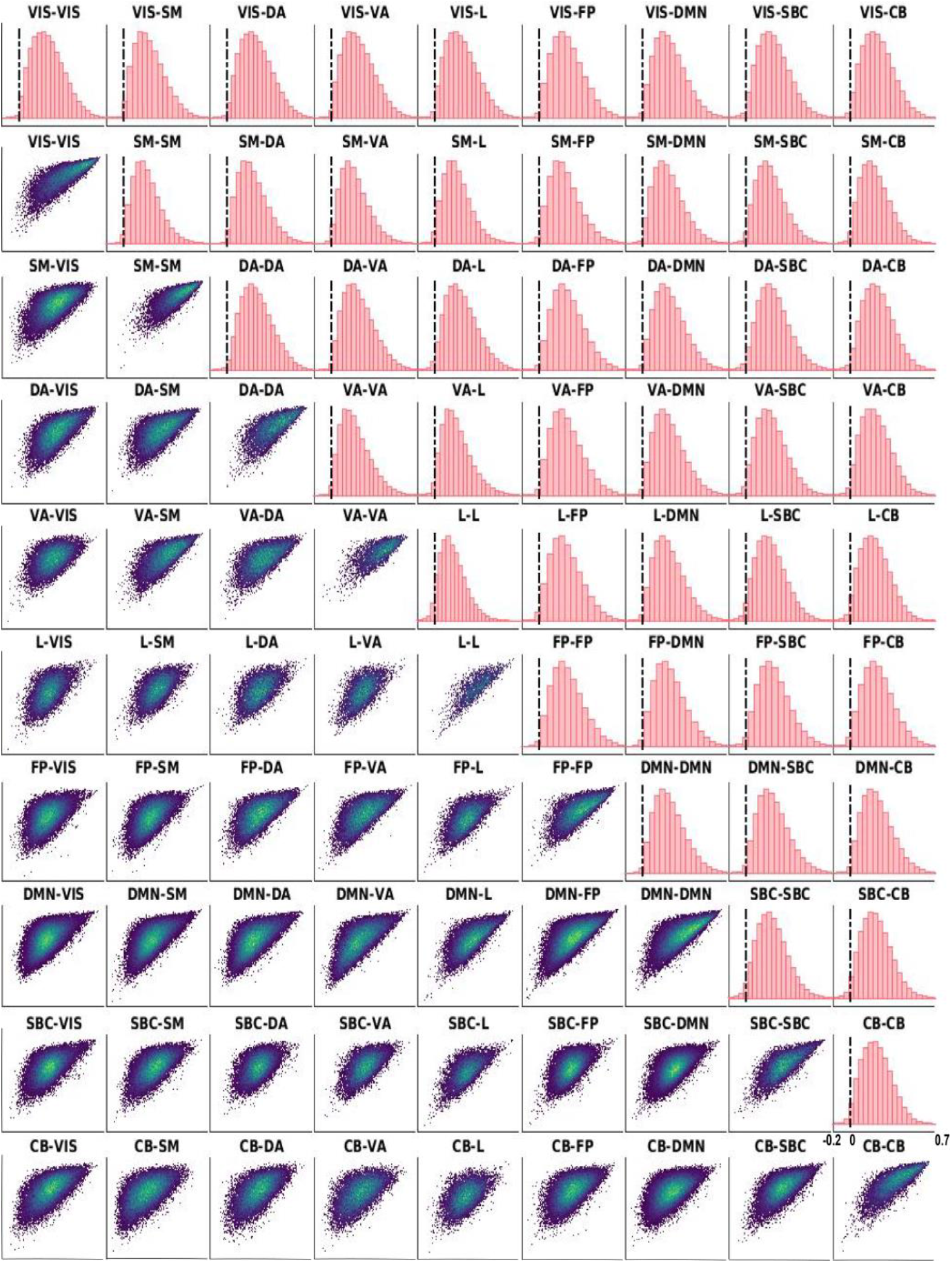
Network-specific effects of GSR on correlational functional connectivity. The top triangle presents the difference histograms between the correlational functional connectivity without and with GSR for each of the ICN pairs. The bottom right histogram shows the scale of correlational differences and is the same for each histogram. The bottom triangle presents the density scattcrplots of the mutual information functional connectivity without GSR (x-axis) and with GSR (y-axis) for each of the ICN pairs. The colormap presents the density from low density (purple) to high density (yellow)

**Figure 5.**
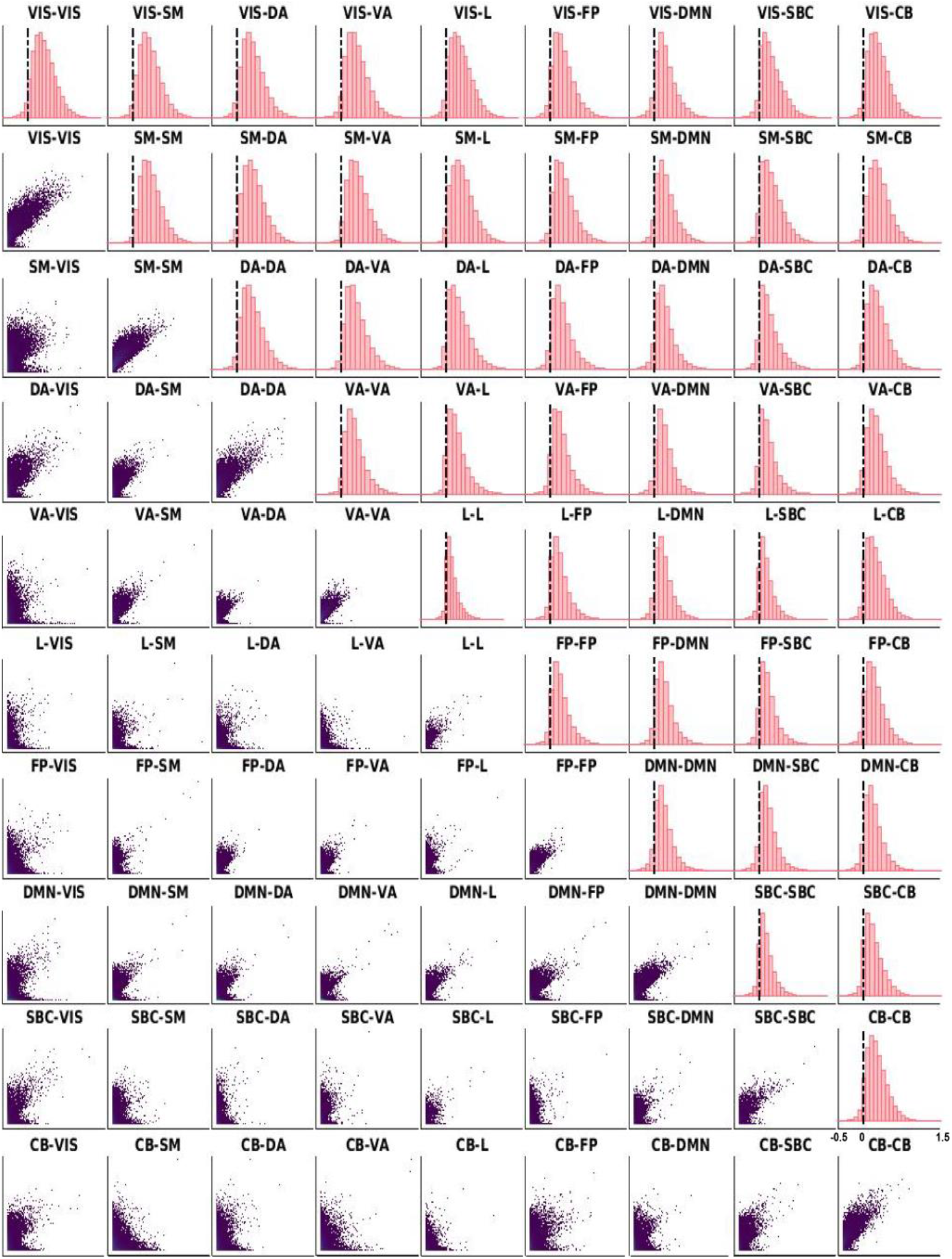
Network-specific effects of GSR on mutual information functional connectivity. The top triangle presents the difference histograms between the mutual information functional connectivity without and with GSR. for each of the ICN pairs. The bottom right histogram shows the scale of mutual information differences and is the same for each histogram. The bottom triangle presents the density scatterplots of the mutual information functional connectivity without GSR (x-axis) and with GSR (y-axis) for each of the ICN pairs. The colormap presents the density from low density (purple) to high density (yellow).

Taken together, GSR reduces the functional connectivity across ROI pairs, and the intrinsic connectivity networks are affected in non-uniform ways. Our findings align with previous work in the literature (Gotts et al., 2013; Saad et al., 2012). Recent work by Li and colleagues who also studied the effect of GSR in a large sample found similar specific regional and network-related effects introduced by GSR (Li, Kong, Liégeois, et al., 2019).

### Relationship of the Global-, vessel BOLD signal and intrinsic connectivity networks

Figure 6 shows the time series of the GS and vessel BOLD signal from an example subject which are highly related to each other. On average, the GS and vessel BOLD signal are strongly and positively correlated (mean correlation coefficient = 0.91, this is significantly different from 0 (t(864) =153, p<0.001). The statistical test was performed after Fisher-z transforming the correlation coefficients.

**Figure 6.**
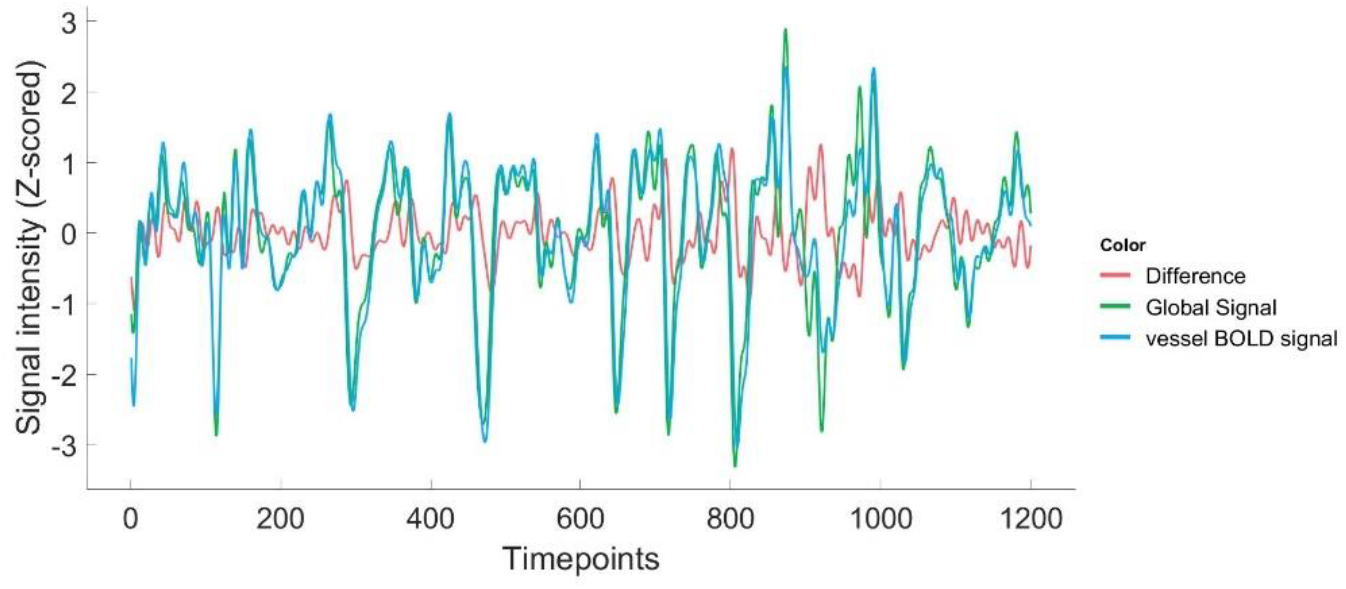
Relationship between the global- and vessel BOLD signal. Example of time series from a subject of the Global signal (green) and vessel BOLD signal (blue) which are highly related to each other. The difference between the global- and vessel BOLD signal is plotted in red.

In Figure 7, the relationship between the GS and the ICN before and after conditioning on the vessel BOLD signal are shown. The GS has an average correlation coefficient of 0.64 and MI of 0.44 with the ICN before controlling for the contribution of vessel BOLD signal. After conditioning on the vessel BOLD signal there is a reduction of the relationship to an average correlation of 0.32 and MI of 0.14. A paired *t*-test shows that this reduction in correlation (t(864) = 109, p<0.001) and MI (t(864) = 71, p<0.001) is significant with large effect sizes for the correlation (Cohen’s đ = 3.72) and MI (Cohen’s d = 2.41).

**Figure 7.**
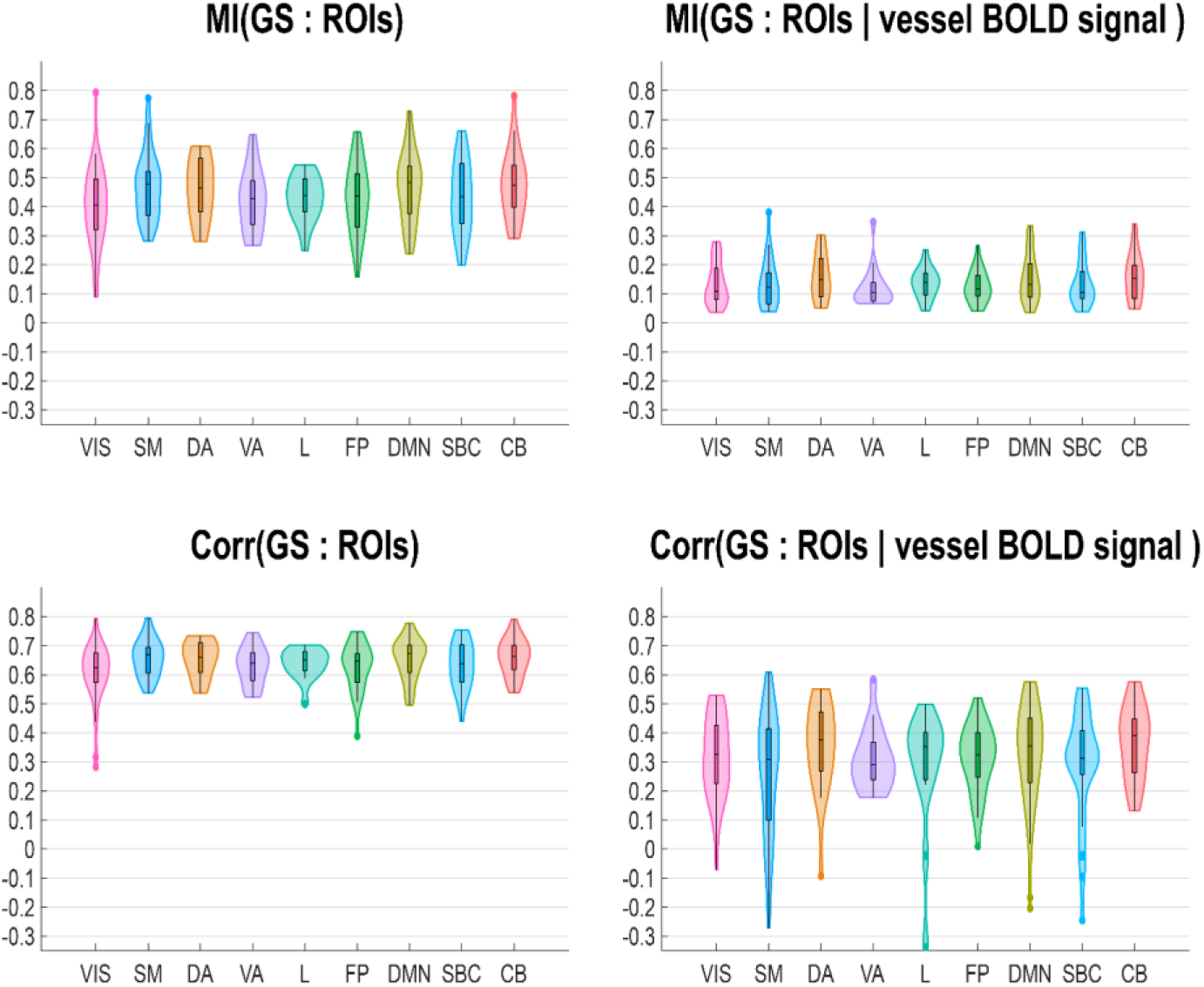
Conditioning role of the vessel BOLD signal. Contribution of the vessel BOLD signal to the relationship between the GS and intrinsic connectivity networks in rs-fMRI data. Top panel: violin plots of the mutual information between the GS and connectome before and after conditioning on the vessel BOLD signal. Bottom panel: violin plots of the correlations between the GS and connectome before and after conditioning on the vessel BOLD signal.

These results indicate that the GS and vessel BOLD signal are highly related to each other. Additionally, we show that the vessel BOLD signal is an important contributing factor to the relationship between the GS and ICN, but does not fully eliminate the relationship. Other groups have found similar results showing that the BOLD signal extracted from arteries and veins are highly correlated with the GS (He et al., 2010; Tong, Yao, Chen, et al., 2018). The sLFO’s that travel with the blood flow through the brain are present in the GS and the vessel BOLD signal, explaining why they arc highly correlated. Despite being highly correlated, our conditioning results on the vessel BOLD signal indicate that both signals have differential and joint effects on the ICNs. However, another framework is needed to further disambiguate the effects of the GS and vessel BOLD signal on the connectome. By applying the PID framework we were able to disambiguate in space and time the differential and joint predictive information for both signals on the different ROIs comprising each ICN.

#### Partial information decomposition

In Figure 8, we depict the terms of the PID, applied from the two drivers (GS and vessel BOLD signal) towards the individual ROIs ordered into the ICNs as a function of timescale. Kruskal-Wallis tests were conducted to examine network-specific effects of information transfer towards the different ICN for each element of the PID. For each element (unique information two sources, synergistic and redundant transfer) the time scale for which the maximum transfer value was attained, and was used to test for differences between the ICNs. For the unique transfer of the GS (Chi square = 2342.7, p < 0.001, df=8), unique transfer of the vessel BOLD signal (Chi square = 2080.36, p < 0.001, df = 8), redundant transfer (Chi square = 455.52, p < 0.001, df = 8) and synergistic transfer (Chi square = 74.03, p < 0.001. df = 8), significant differences were found among the nine ICNs (VIS, SM, DA, VA, L, FP, DMN, SBC, CB). These tests show that for each of the PID terms, the information transfer towards the ICNs are different at the time scale of maximum information transfer, confirming network-specific effects.

**Figure 8.**
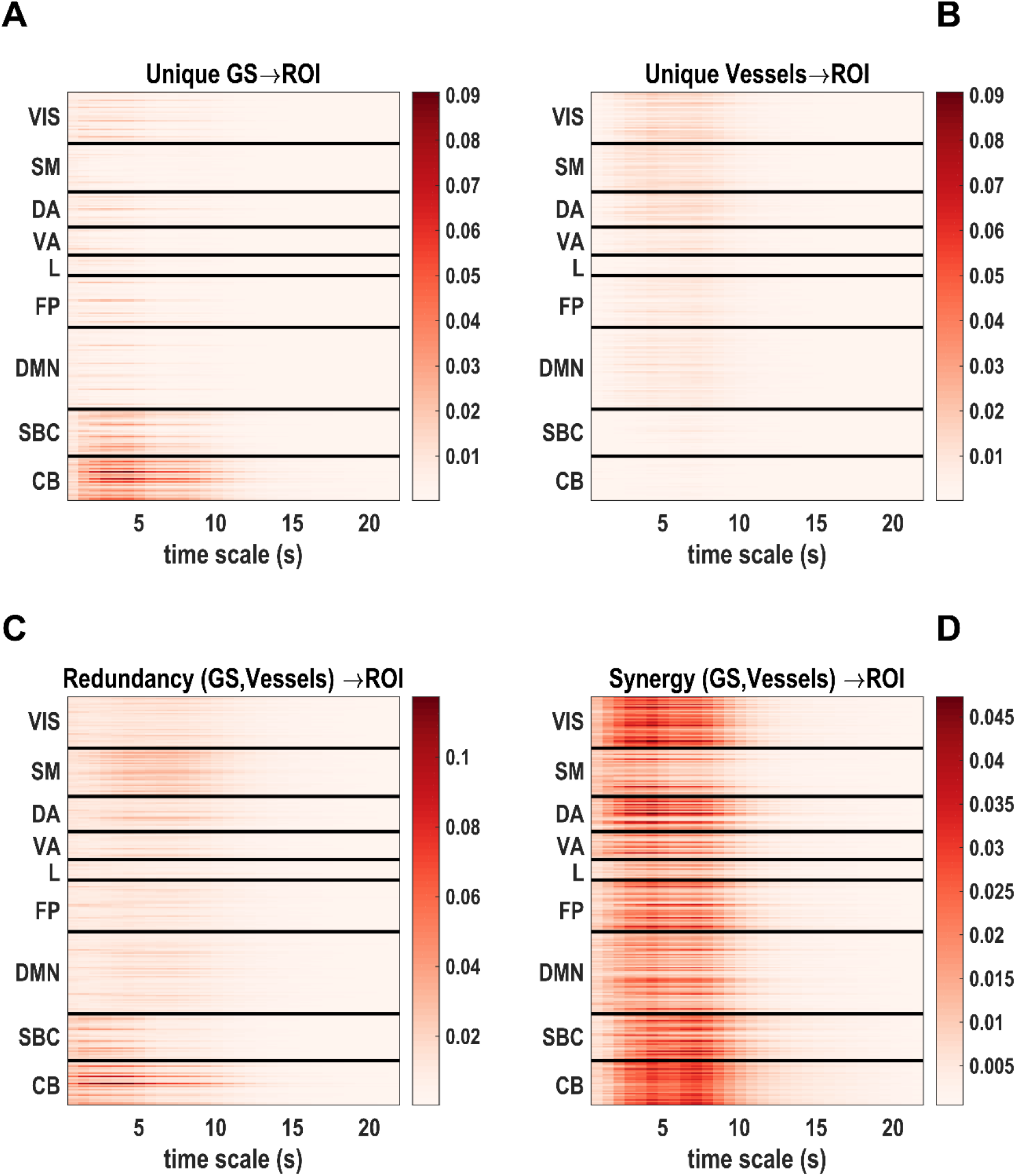
Partial information decomposition (PID) in the HCP rs-fMRI data. Partial information decomposition (PID) of the information transfer from the GS and vessel BOLD signal to the individual ROIs. ***(A):*** the unique information from the GS to the individual ROIs; ***(B):*** the unique information from the vessel BOLD signal to the individual ROIs. ***(C):*** the redundant transfer from the GS and vessel BOLD signal to the individual ROIs. ***(D):*** the synergistic transfer from the GS and vessel BOLD signal to the individual ROIs.

Furthermore, as depicted in Figure 9, the significant differences between the unique information from the CS and vessel BOLD signal is different towards the ICNs across timescales, assessed with a permutation based paired *t*-test. Interestingly, the BOLD signal from the vessels provide more unique information to the cortical ROIs at a time scale corresponding to the one of the hemodynamic response function, peaking on average around 6-7 seconds. The unique GS information has high levels of transfer towards the CB and SBC (Figure 8A). The unique vessel BOLD signal information has high levels of transfer towards the VIS, SM, DA, VA, FP and DMN (Figure 8B). It’s clear from the PID that both synergistic and redundant transfer occur at the same time across ICNs and multiple time scales (Figure 8C, D).

**Figure 9.**
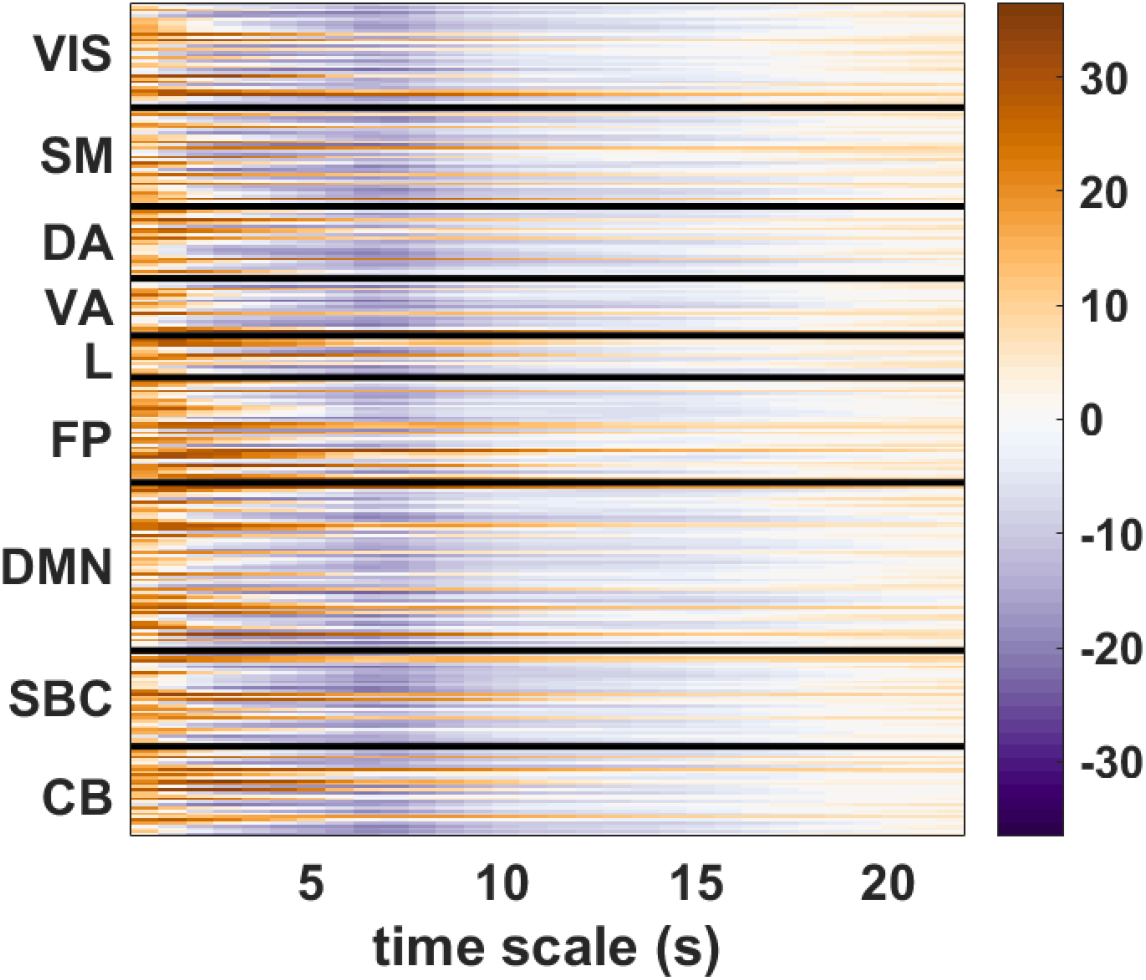
Statistical significance between Unique GS and unique vessel BOLD signal. paired *t*-test using permutations between the unique GS and unique vessel BOLD signal for each ROI at each timescale. Warm colors (positive values) indicate that the GS provides more unique information to the cortical ROIs than vessel BOLD signal, while cold colors (negative values) indicate a greater unique contribution of vessel BOLD signal.

We observe by applying PID that the information terms are modulated across two different domains. 1) Spatial modulation: both the GS and vessel BOLD signal despite being highly related, affect the ICNs in different ways as evidenced by the difference in information transfer of the unique information transfer’s. At the same time, the joint information (dynamics) of both signals is also important as they modulate the ICNs in terms of synergistic and redundant transfer. This spatial modulation of information transfer is confirmed by the Kruskal-Wallis tests; 2) Temporal modulation: as evidenced by all terms, transfer happens at multiple timescales and patterns of transfer towards ICNs can change depending on the timescale. Even more, the importance of decomposing information at multiple scales is testified by the fact that terms (e.g. unique GS, unique vessel blood, synergy and redundancy …) attain maximum transfer values at time scales > 1.

### Simulations

In the simulations we attempt to explain the spatial and temporal modulations of the information decomposition. We hypothesized that blood arrival time might be driving the observed spatiotemporal modulations. In the simulations a common ‘blood component’ carrying physiological information (sLFO’s) was mixed with interacting neuronal components and arrived with varying times at the components.

#### GSR

The results of applying GSR on the simulated interacting rs-fMRI components are depicted in Figure 10. Figure 10A shows the functional connectivity using correlations among the 9 simulated components, averaged across 100 simulations without (bottom triangle) and with GSR (top triangle). Figure 10B shows the functional connectivity using mutual information for the same dataset with and without GSR. In the correlational approach, GSR reduces functional connectivity and introduces anti-correlations across component pairs. The same observations are made in the mutual information approach, where GSR reduces the functional connectivity across component pairs. In both cases, we observe that the reduction in functional connectivity follows a gradient across component pairs from component 1 to 9.

**Figure 10.**
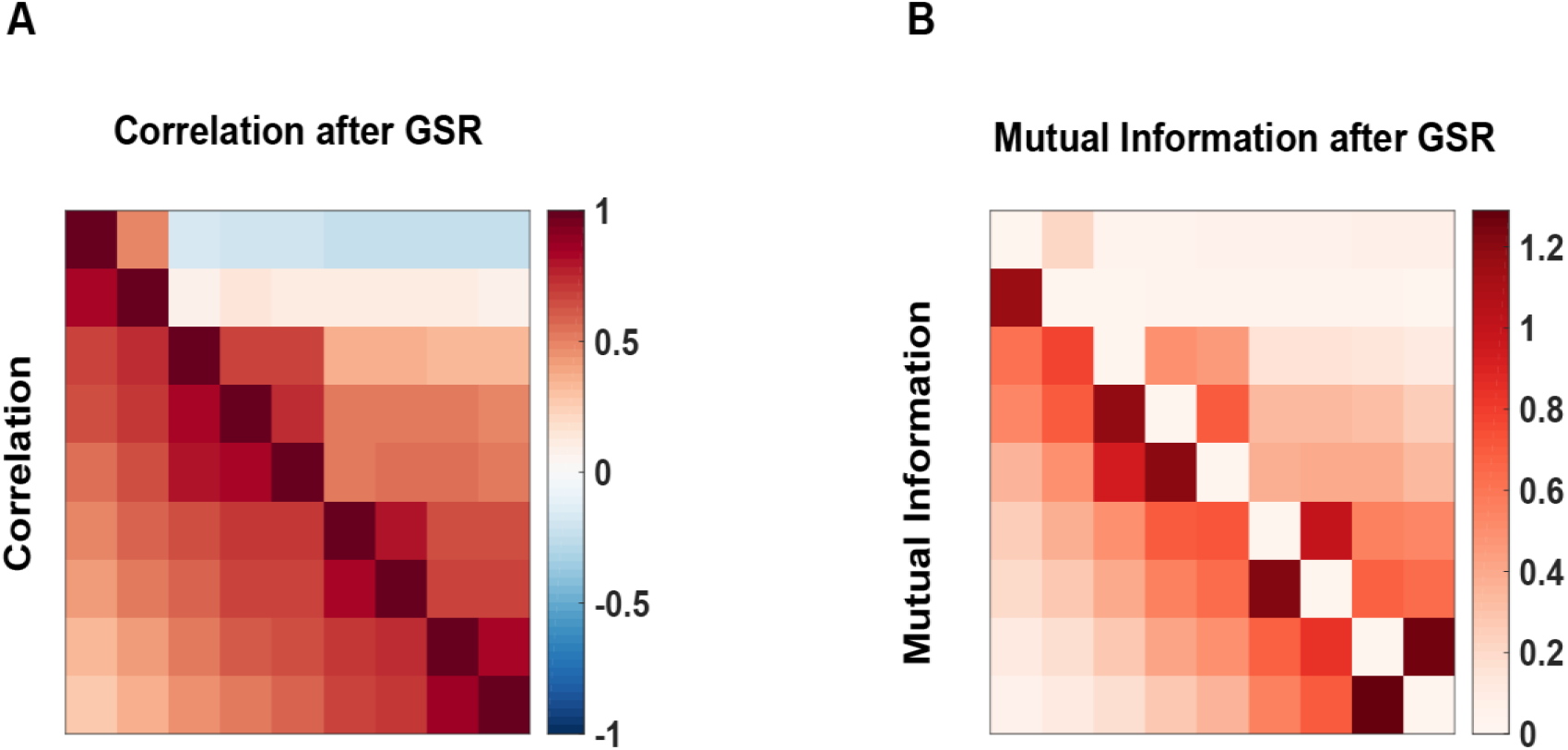
Effect of GSR on functional connectivity in the simulated dataset. ***(A)*** estimated functional connectivity using correlations before (bottom triangle) and after GSR (top triangle). ***(B)*** estimated functional connectivity using mutual information before (bottom triangle) and after GSR (top triangle). Interacting rs-fMRI components were simulated (1 to 9). Blood arrival time (BAT) was linearly increased from component 1 to 9. In component 1 BAT is earlier compared to late BAT in component 9.

This gradient can be explained by the difference in blood arrival time (BAT) between components, that was linearly increased during the simulations. Similar as in our empirical rs-fMRI HCP results, where GSR has network-specific effects across regions and ICNs, we observe similar results in our simulations.

#### Partial Information Decomposition

In Figure 11, we depict the terms of the PID, applied from the two drivers (GS and vessel BOLD signal) towards the simulated components ordered according to BAT as a function of timescale. For each of the PID terms depicted in Figure 11A (unique information GS), Figure 11B (unique information vessel BOLD signal), Figure 11C (redundancy) and Figure 11D (synergy), we observe a gradient in the information transfer patterns that can be explained by the BAT. Figure 11 A, shows the unique information from the GS towards the components. In this case, maximum transfer is attained in components with an early BAT. Components with an early BAT show high information transfer across low timescales, while at higher timescales the transfer reduces gradually. Components with late BAT show almost no transfer across low timescales and some transfer at higher timescales.

**Figure 11.**
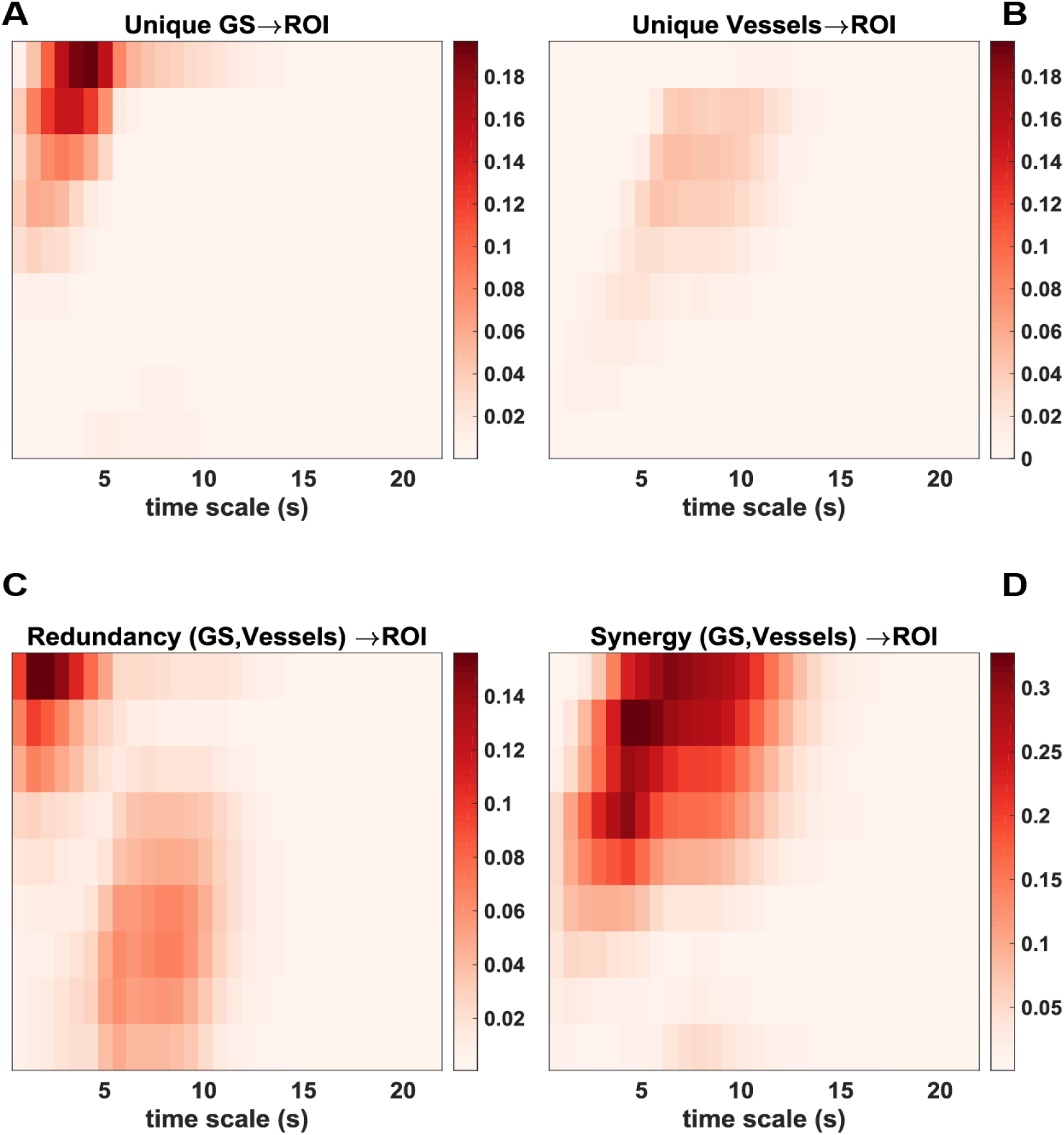
Partial information decomposition (PID) in simulated rs-fMRI data. Partial information decomposition (PID) in simulated rs-fMRI data of the information transfer from the GS and vessel BOLD signal towards the individual ROIs. ***(A)*** the unique information from the GS to the individual ROIs. ***(B)*** the unique information from the vessel BOLD signal to the individual ROIs. ***(C)*** the redundant transfer from the GS and vessel BOLD signal to the individual ROIs. ***(D)*** the synergistic transfer from the GS and vessel BOLD signal to the individual ROIs.

Figure 11B shows the unique information from the vessel BOLD signal towards the components. With a linearly increasing BAT, information transfer is observed at lower timescales. Here it is clear from the patterns of information transfer observed in Figure 11A and Figure 11B that both drivers (GS and vessel BOLD) effect the simulated components in a differential way. However, at the same time both signals show joint dynamics of information transfer towards the simulated components. This is clear from Figure 11C and 11D, which show the redundant and synergistic transfer, respectively. In Figure 11C, the maximum redundant transfer is attained in components with an early BAT. Components with an early BAT show high information transfer at lower timescales, and this transfer decreases with increasing timescales. On the other hand, components with a late BAT show an increase in transfer from lower to higher timescales. Figure 11D shows the synergistic transfer of information. Components with an early BAT show high transfer of information across low timescales and this decreases with increasing timescales. Components with a late BAT show lower transfer across all timescales.

The results from the PID observed from the simulations are similar to the PID results from the empirical HCP rs-fMRI dataset (Figure 8). With the simulations the goal was to explain the spatiotcmporal modulations observed in Figure 8 by varying the BAT in interacting components. Indeed, in Figure 11, we observe that with our simulations we observe similar spatiotemporal modulations of the PID terms. The difference between both modalities is that in the simulations we were able to order our components according to BAT (Figure 11), while the rs-fMRI HCP data is ordered in ICNs and not according to BAT (Figure 8).

### Simultaneous calcium and hemodynamic recordings

#### GSR

Figure 12A shows the functional connectivity using correlations among the 38 ROIs without (bottom triangle) and with GSR (top triangle) in the single mouse hemodynamic recordings. Figure 12B shows the functional connectivity using mutual information for the same dataset with and without GSR. In the correlational approach, GSR reduces functional connectivity across almost all ROI pairs. Anti-correlations are introduced across ROIs in a non-uniform way across the connectome. The same observations are made in the mutual information approach, where GSR reduces the functional connectivity across ROI pairs. These findings are similar to the results we found with the HCP rs-fMRI and simulated rs-fMRI dataset (Figure 3, 10). Namely, GSR, has regional and network-specific effects in reducing functional connectivity.

**Figure 12.**
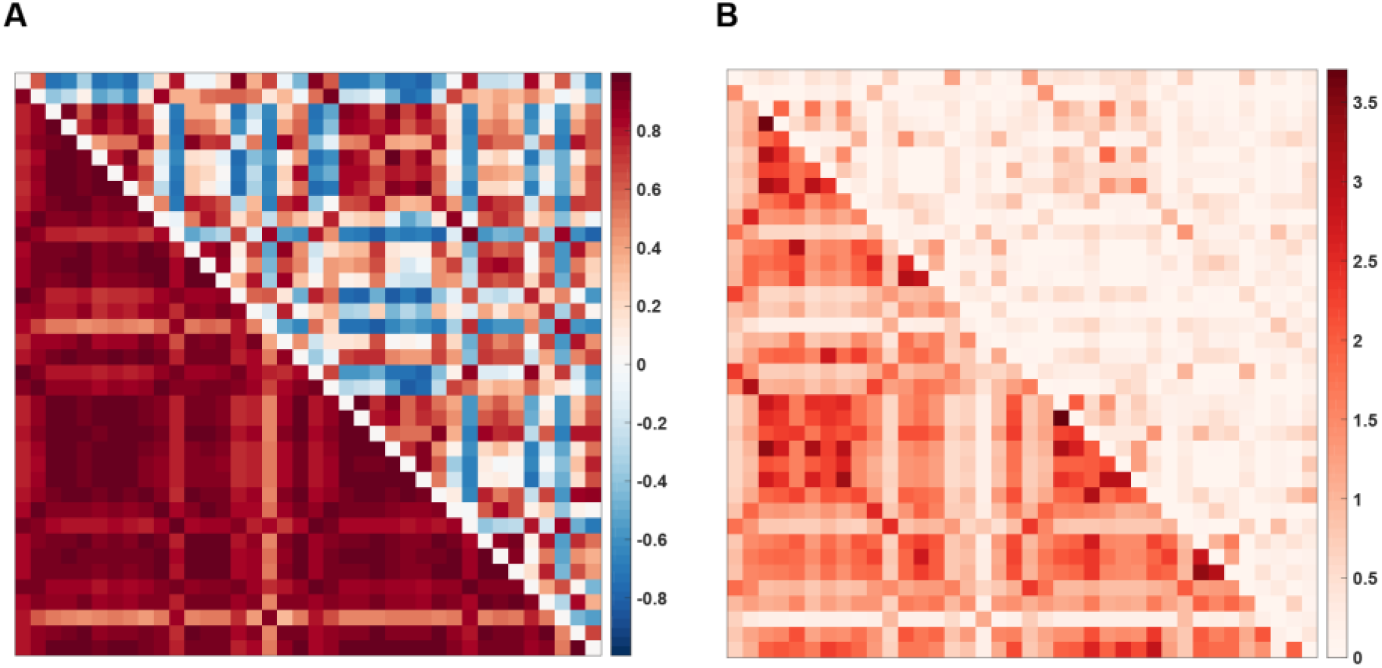
Effect of GSR on functional connectivity in hemodynamic mouse recordings. ***(A)*** estimated functional connectivity using correlations before (bottom triangle) and after GSR (top triangle). ***(B)*** estimated functional connectivity using mutual information before (bottom triangle) and after GSR (top triangle). The same 38 ROIs were used as described in (Matsui et al., 2016, 2018b).

#### Partial Information Decomposition

In Figure 13 A-D, we depict the results of the PID applied to the hemodynamic recordings. The terms of the PID, applied from the two drivers (homodynamic GS. hemodynamic vessel BOLD signal) towards the individual hemodynamic ROI signals as a function of timescale are shown. For each of the PID components, we observe a spatial and temporal modulation of information transfer towards the individual RO1 signals of the mouse. From the unique information from the GS and vessel BOLD signal (Figure 13A, B) we observe transfer that is different towards the individual ROIs and timescales. Furthermore, both signals show joint transfer of redundant and synergistic information (Figure 13C, D) which is different for the ROIs across lower and higher timescales.

**Figure 13.**
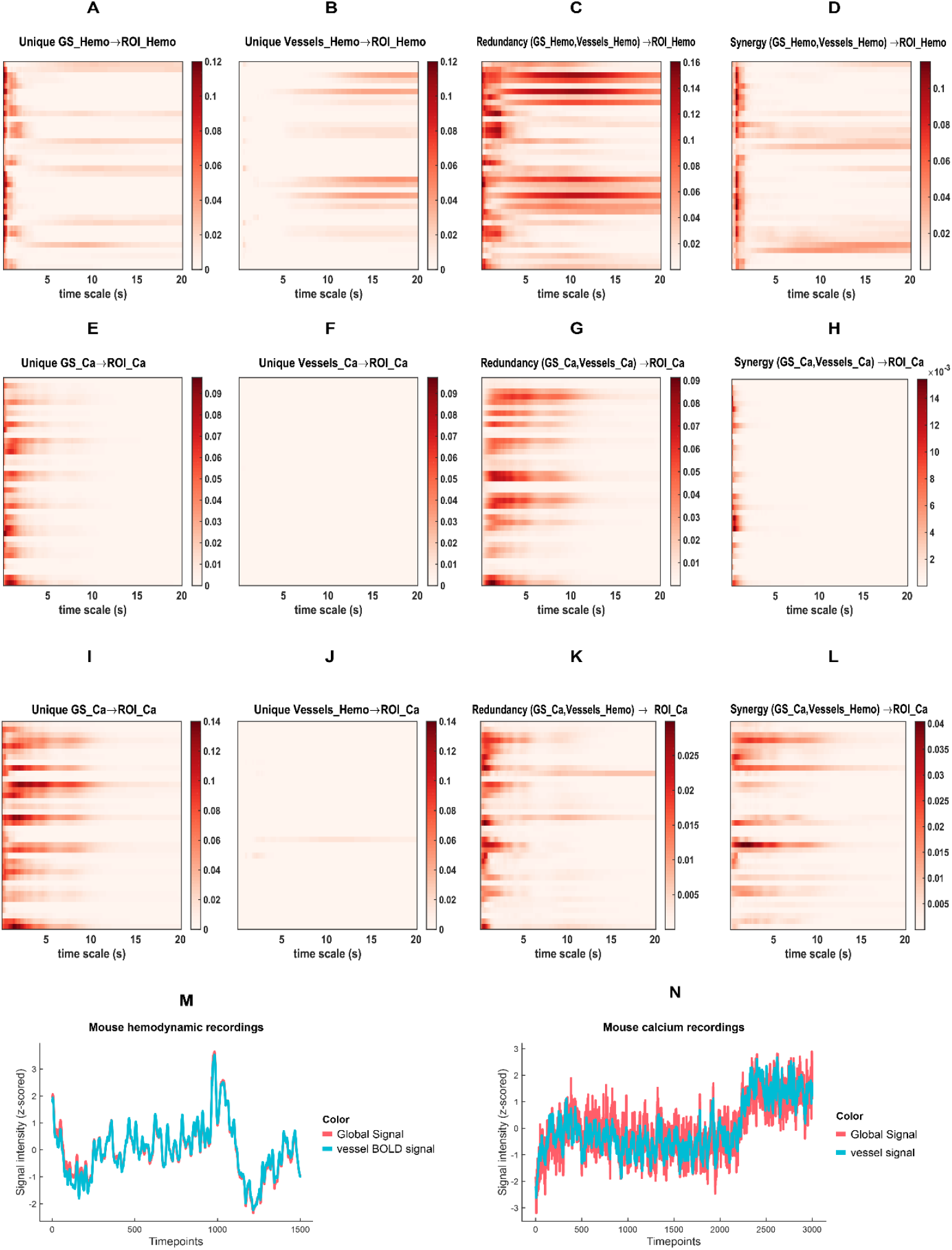
Partial Information Decomposition (PID) in hemodynamic and calcium mouse recordings. PID in hemodynamic mouse recordings of the information transfer from the GS and vessel BOLD signal towards the individual ROIs. ***(A)*** the unique information from the hemodynamic GS to the individual hemodynamic ROI signals ***(B)*** the unique information from the hemodynamic vessel BOLD signal to the individual hemodynamic ROI signals. ***(C)*** the redundant transfer from the hemodynamic GS and vessel BOLD signal to the individual hemodynamic ROI signals. ***(D)*** the synergistic transfer from the hemodynamic GS and vessel BOLD signal to the individual hemodynamic ROI signals. ***(E)*** the unique information from the calcium GS to the individual calcium ROI signals. ***(F)*** the unique information from the calcium vessel signal to the individual calcium ROT signals. ***(G)*** the redundant transfer from the calcium GS and vessel signal to the individual calcium ROI signals. ***(H)*** the synergistic transfer from the calcium GS and vessel signal to the individual calcium ROI signals. ***(I)*** the unique information from the calcium GS to the individual calcium ROI signals ***(J)*** the unique information from the hemodynamic vessel BOLD signal to the individual calcium ROI signals. ***(K)*** the redundant transfer from the calcium GS and hemodynamic vessel BOLD signal to the individual calcium ROI signals. ***(L)*** the synergistic transfer from the calcium GS and hemodynamic vessel BOLD signal to the individual calcium ROI signals. ***(M)*** time series of the GS (red) and vessel BOLD signal (blue) in hemodynamic recordings, which are highly correlated (0.99). ***(N)*** time series of the GS (red) and vessel signal (blue) in calcium recordings which are also highly correlated (0.89).

In Figure 13 E-H, we depict the results of the PID applied to the calcium recordings. The terms of the PID, applied from the two drivers (calcium GS, calcium vessel signal) towards the individual calcium ROI signals as a function of timescale are shown. In this case, only a spatial and temporal modulation of transfer is observed in the unique information from the GS (Figure 13E) and redundant information (Figure 13**Error! Reference source not found.**G). The transfer of unique information from the vessel BOLD signal observed in the hemodynamic recordings (Figure 13B) disappears in calcium recordings (Figure 13E), as well as most of its synergy with GS, except from short time scales (Figure 13H). This result should be expected as calcium recordings reflect more prominently neuronal activity in contrast to hemodynamic recordings, therefore the calcium vessel signal is expected not to contain physiological information. As a result: 1) there is no unique information from calcium vessel signal (Figure 13E) and synergistic transfer compared (Figure 13H) with the hemodynamic recordings (Figure 13B, D), where the hemodynamic vessel BOLD signal does contain physiological information; 2) The pattern of redundant transfer (Figure 13G) can then most likely be explained by noise in the calcium vessel signal, which shares some component with the neuronal GS and predicts the information towards the individual ROIs. One possible source of noise could be from the optical imaging method. Some fluctuations could be induced in the signals by the intensity of excitation light which could affect the entire field of view if it exists. The pattern of redundant transfer in the hemodynamic recordings (Figure 13C) is due to the sharing of physiological information in the GS and vessel BOLD signal.

In Figure 13 I-L, we depict the results of the PID applied to a combination of the hemodynamic and calcium recordings, after downsampling the latter by a factor 2 to match the former. The terms of the PID, applied from the two drivers (calcium GS, hemodynamic vessel BOLD signal) towards the individual calcium ROI signals as a function of timescale are shown.

In this case, we observe a spatial and temporal modulation of transfer in the unique information from the GS (Figure 13I) and redundant and synergistic information (Figure 13**Error! Reference source not found. K**, L). Here the effect of unique information from the hemodynamic vessel BOLD signal disappears (Figure 13J). This shows us that there is no transfer of information of hemodynamic vessel BOLD signal towards calcium ROI signals. This gives us evidence that neuronal functional activity is unaffected by the fluctuations of hemodynamic vessel BOLD signal.

## Discussion

In this paper we learnt that, together with their very high correlation, the global signal and the BOLD signals in the vessels provide both unique and shared contributions to the reduction in uncertainty on the dynamics of cortical areas. These differential actions contribute to explaining the effect of GSR on brain-wide intrinsic functional connectivity. Additionally, the presence of unique information from both signals could indicate that it is possible, at least theoretically, to separate the two types of signals, hopefully keeping the neural and cognitive correlates, while discarding the physiological nuisance.

Our observations after applying GSR to a large rs-fMRI dataset are consistent with the literature. Using a correlational and mutual information framework, we observed that GSR reduces the functional connectivity between ROI pairs (Figure 3A, B). This is in line with previous work (M. D. Fox, Zhang, Snyder, & Raichle, 2009; Murphy et al., 2009; Wcisscnbachcr et al., 2009). Moreover, we observed network-specific effects. The reduction in functional connectivity was not uniform across regions and networks (Figure 4, 5). This is consistent with previous findings from (Gotts et al., 2013; Saad et al., 2012), and recent work by (Li, Kong, Liégeois, et al., 2019) who also studied the effect of GSR in large human rs-fMRI datasets. These findings were verified in the simulations (Figure 10) and cross-species mouse recordings (Figure 12).

In a next step, we investigated the role of the global signal and vessel BOLD signal in this modulation and further mapped their presence across intrinsic connectivity networks and time using PID. First, we found that the GS and vessel BOLD signal are highly related (Figure 6), as reported in previous work (Tong, Yao, Jean Chen, et al., 2018). According to Tong and colleagues this finding can be explained by the fact that slow low frequency oscillations (sLFO’s) are widespread across the brain travelling with the cerebral blood flow. During the calculation of the GS, neuronal activation which is more local is smoothed/cancelled out leaving the sLFO’s as the dominant signal in both the GS and vessel BOLD signal. We further found that the vessel BOLD signal is an important contributor between the relationship of the GS and the intrinsic connectivity networks, but does not fully explain the relationship (Figure 7). So, despite being highly related they both seem to have some unique relatedness to the intrinsic connectivity networks. By applying PID, we were able to disambiguate the unique and joint dynamics of the GS and vessel BOLD signal by quantifying their presence across networks and time.

We observed in empirical rs-fMRI data, simulations, and hemodynamic mouse recordings that the predictive information from the GS and vessel BOLD signal is present in different amounts across time and space (Figure 8, 11, 13). The information of the GS and vessel BOLD signal are present across the connectome in unique (Figure 8A, B; 11 A, B; 13A, B) and joint (Figure 8C, D; 11C, D; 13C, D) ways. Furthermore, we confirmed that: 1) the spatiotemporal modulation of the predictive information in human rs-fMRI (Figure 8) can be explained in terms of blood arrival time in the simulations (Figure 11); and 2) the spatiotemporal modulation is due to the presence of physiological information (sLFO’s) in hemodynamic signals in the vessel BOLD signal (Figure 8B, 13B). As a result, the predictive information of the vessel BOLD signal disappears in calcium recordings due to the lack of physiological information in neuronal signals (**Error! Reference source not found.F**).

Taken together, without applying GSR both signals are present in different amounts and with different timing across regions/ networks and one should be aware of this when interpreting functional connectivity estimates. We know that the GS is related to physiology (vascular component), and this is due to non-neuronal sLFO’s travelling with the blood flow (Tong, Yao, Chen, et al., 2018). For denoising strategies that aim to remove non-neuronal artifacts such as motion or-, respiration, it could be beneficial to control for the sLFO’s by correcting for blood arrival time.

The classic GSR approach has been shown to be effective in accounting for global motion and respiratory artifacts compared with other de-noising approaches (Ciric et al., 2017; Lydon-Staley et al., 2018; Parkes et al., 2018; Power, Plitt, et al., 2017; Satterthwaite et al., 2017). However, it does not consider the dynamic passage of global systemic effects (sLFO’s) throughout the brain. The dynamic GSR method by (Erdoğan et al., 2016) can effectively deal with the sLFO’s by correcting for blood arrival time and reduces the impact of physiological noise. As a result, the functional connectivity estimates provide enhanced specificity and reflect more the underlying biology rather than spurious noise. However, it is not known if the dynamic GSR, method removes all physiological artifacts. It controls for the vascular effect of the sLFO’s, but other physiological effect such as Mayer waves that are related to the sympathetic nervous system oscillations, might be another potential source of confounding global signal oscillations (Julien, 2006).

From the work of Saad and colleagues we know that GSR alters the underlying correlational structure in unpredictable ways depending on the contribution of regions/networks to the GS (Saad et al., 2012). From our observations we have shown that physiology is related to the GS and present in different amounts across networks. Following this view, regressing out the GS might selectively introduce new-mixed physiological artifacts in the functional connectivity estimates across networks if not controlled for. As a result, these new mixed physiological artifacts could confound the interpretations made from functional connectivity estimates. Being aware of this is important as a wave of recent studies have shown great interest in the relationship between behavior and the architecture of the brain uncovered by functional connectivity (Dubois & Adolphs, 2016; Finn et al., 2015; Kong et al., 2018; Li, Kong, Liégeois, et al., 2019; Rosenberg et al., 2015). Recent work by Li and colleagues shows that GSR strengthens the association between functional connectivity and behavioral measures. One could argue that the reliable association between functional connectivity could be partly explained by the introduced physiological artifacts after GSR. Future work that focuses on the relationship between behavior and connectivity could investigate if controlling for the physiology (sLFO’s) by blood arrival time improves the association between behavior and functional connectivity even more.

While global fluctuations have been linked to artifacts such as participant motion, respiration and sLFO’s, other studies have linked global fluctuations to neuronal origins. Fluctuations of the rs-fMRI GS have been linked to vigilance (Wong et al., 2013), glucose metabolism (Thompson et al., 2016) and arousal mediated by ascending nuclei (X. Liu et al., 2018; Turchi et al., 2018). More recently Gutierrez-Barragan and colleagues have shown that each cycle phase of the GS is the sum of differently overlapping recurring brain states that have a neuronal origin (Gutierrez-Barragan, Basson, Panzeri, & Gozzi. 2018). These brain states that govern the spontaneous brain activity can be captured by CAP-based approaches (Gutierrez-Barragan et al., 2018; Karahanoğlu & Van De Ville, 2015; X. Liu, Chang, & Duyn, 2013; X. Liu & Duyn, 2013; Matsui et al., 2016). In addition, work examining global signal topography demonstrates structured information in the GS that can explain individual differences in trait-level cognition and behavior, further suggesting that the signal contains strong cognitive influences (Bolt, Li, Bzdok, Nomi, Yeo, Spreng, Uddin, under review). Aside from the introduction of mixed physiological artifacts, we do not know the additional effects of GSR on recurring brain states with a neuronal origin and how this is reflected in functional connectivity estimates.

To conclude, the current study is not meant to suggest a particular stance regarding the use of GSR in studies of rs-fMRI functional connectivity. Rather, the goal is to highlight the finding that the GS and physiology are related and present in different amounts across networks. If one wants to apply de-noising strategies such as GSR, one should be aware of the physiological confound in functional connectivity estimates. An alternative de-noising strategy that could be considered is dynamic GSR, which effectively reduces the impact of physiological artifacts in functional connectivity patterns (Erdoĝan et al., 2016). On the other hand, we know that GSR also impacts task-relevant neuronal information, further complicating the interpretation of functional connectivity findings. Alternative de-noising approaches that retain neuronal information and remove global artifacts based on temporal ICA might be promising alternatives to GSR (Glasser et al., 2018, but see also the comment by Power, 2019).

### Limitations

To extract the vessel BOLD signal from the human rs-fMRI data, we created masks that contained vessel information after thresholding the T1w/T2w ratio images. However, we realize that due to the low spatial resolution of tMRI our masks can contain voxels that could be classified as grey matter voxels. As a result, the BOLD signal of our vessel BOLD signal mask might be a mixture of grey matter- and vessel BOLD signal effects. Despite the possibility of mixing effects in the vessel BOLD signal that could bias our results, we believe the findings using PID to be robust (Figure 8, 13):

1. Using mouse data, we found a spatiotemporal modulation of the predictive information just as in human rs-fMRI data (Figure 8, 13). Compared with human rs-fMRI data, the mouse data has the advantage of having high spatial resolution, and reduced movement since the animal is constrained in a rig. Due to the higher resolution we were able to create a mask that only contains vessel information. The extracted vessel BOLD signal in this case is not a mixture of neuronal and vessel information. As the findings are similar as in human data (including a very high correlation of hemodynamic signals in the vessels and global signal, Figure 13M, N), we are confident that a possible mixing of the vessel BOLD signal is not confounding the current results. Further, we found that the unique predictive information of the vessel BOLD signal is absent in calcium recordings (Figure 13E) and was only found with hemodynamic recordings in mouse data (Figure 13B). This is because calcium vessel signal does not contain physiological (sLFO’s) information.
2. In PID, the measures explicitly look for contributions above and beyond what is contained in the target, meaning that the simple co-existence of two phenomena in a time series does not count in the PID.
3. In the human rs-fMRI we used a large sample, averaging the results over hundred of subjects. We could expect that the location of grey matter voxels and neuronal information that might contribute to the vessel BOLD signal mask is inconsistent across subjects. Therefore, the expected neuronal mixing effect might be averaged out in the large group analysis we performed. On the other hand, it is still possible that a neuronal contribution to the vessel BOLD signal is present across subjects and potentially biases the results even in a large sample size.

To summarize, even though what we call vessel BOLD signal in this work (human rs-fMRI) might be a mixture of neuronal and vessel information, we are quite confident that the observed spatiotemporal modulation of the PID in human rs-fMRI is due to the physiological information (sLFO’s) that is present in the vessel BOLD signal. Work that aims to validate the results in human rs-fMRI, excluding any neuronal contamination effects, might consider extracting the mask for the vessel BOLD signal from subjects that have angiography data available.

## Conclusion

We confirmed that GSR reduces functional connectivity estimates between regions and networks in a large empirical fMRI dataset. Furthermore, using PID we show that the GS and sLFO’s (physiological artifact) are present in different amounts across different timings, regions and networks. Using simulations we were able to explain the spatiotemporal modulation of the PID in terms of blood arrival time. Thus, correcting for the sLFO’s by taking blood arrival time into account might reduce the introduction of physiological artifacts in functional connectivity.

## Acknowledgments

We thank Teppei Matsui for sharing the mouse data, and for comments that improved the manuscript.

This research was supported by the Fund for Scientific Research-Flanders (FWO-V, grant FWO16-ASP-050 awarded to NC). The computational resources (Stevin Supercomputer Infrastructure) and services used in this work were provided by the VSC (Flemish Supercomputer Center), funded by Ghent University, FWO and the Flemish Government - department EWI.

